# Refractive index as an indicator for dynamic protein condensation in cell nuclei

**DOI:** 10.1101/2025.04.18.647244

**Authors:** Orlando Marin, Peter Kirchweger, Arina Dalaloyan, Yoav Barak, Michael Elbaum

## Abstract

Protein condensation is the basis for formation of membrane-less organelles in the cell. Most famously, weak, polyvalent interactions, often including RNA, may lead to a liquid-liquid phase separation. This effect greatly enhances local concentrations and is thought to promote interactions that would remain rare in dilute solution. Synthetic systems provide a means to clarify the underlying biophysical mechanisms at play, both in vitro and in the cell via exogenous expression. In this regard, ferritin is a useful substrate, as its composition of 24 subunits with octahedral symmetry supports self-assembly by close packing in 3D. The conventional diagnostic tool for protein condensation is fluorescence imaging. In this work we explore the use of refractive index mapping to detect states of condensation and decondensation. Using two related ferritin-based self-assembly systems, we find that refractive index is a sensitive indicator for reversible condensation. Surprisingly, refractive index indicates a rapid decondensation even when molecular dispersal kinetics are slow according to fluorescence. Conversely, in a photo-activated condensation where long activation results in slow decondensation kinetics, the refractive index provides reliable evidence for the physical state independent of fluorescence. The observations suggest a distinction between condensation to a sparse biomolecular network, or to a material continuum that supports an optical polarizability distinct from that of the dilute phase in solution.

## Introduction

Macromolecular condensation is recognized as an important mechanism in cellular biochemistry. Interaction between proteins and/or nucleic acids, whose average concentration is low in the cell, can be accelerated and regulated if brought into proximity by co-condensation into a common dense phase. This thermodynamic phase separation underlies the concept of membrane-less organelles, and the dense regions are commonly known as phase condensates (Banani et al., 2017). Their state is often fluid (Brangwynne et al., 2009), leading to the description as liquid-liquid phase separation. The fluid state facilitates molecular mixing and interaction within the condensate.

Mechanistically, phase condensation reflects the effect of weak but polyvalent interactions between binding partners. Many proteins are recognized for their marginal stability in solution, often leading to aggregation or organized self-assembly (Garcia-Seisdedos et al., 2017; Schweke et al., 2024). This may involve a drastic structural rearrangement, for example, in amyloid formation (Sawaya et al., 2021), or in ordered self-assembly as occurs in cytoskeletal or flagellar filaments. Intrinsically disordered protein (IDP) domains are particularly prone to condensation due to their weak and transient hydrophobic interactions, as well as to amyloid-like misfolding (Mukherjee et al., 2024). Indeed, misfolding of common IDPs is a hallmark of many neurodegenerative diseases (Elbaum-Garfinkle, 2019). RNA binding is a common motif for promoting heterogeneous interaction and condensation; the disordered nucleic acid provides a flexible substrate for protein interaction. On the other hand, phase separation can be viewed more strictly as a self-assembly phenomenon. For example, extended supramolecular structures can be constructed from designed peptides of chosen shapes and forms (Heidenreich et al., 2020).

In a similar spirit, self-assembly of a hybrid ferritin fusion to self-dimerizing fluorescent proteins (FPs) led to the formation of spherical bodies (Bellapadrona and Elbaum, 2014). These supramolecular protein assemblies (SMPA) form spontaneously in living tissue culture cells upon expression of the constituents from a transfected plasmid. Assembly could be directed to the cell nucleus by inclusion of a nuclear localization signal (NLS) sequence to the FP. The symmetry of the ferritin core with 24 subunits provides a convenient building block for self-assembly because a sphere in a close-packed pile has 12 nearest neighbors; therefore, weak interactions at the molecular level can induce self-assembly at a much larger scale. When formed in the cell nucleus, evidence of a crystalline structure was observed. The overall structure was often hollow or aveolar, suggesting a sintering of smaller sub-assemblies, but not a liquid-like state. Formed in bacteria, on the other hand, the assemblies showed no long-range molecular order (Bellapadrona et al., 2015). Self-assembly depends on antiparallel dimerization at a hydrophobic patch of amino acids common to green fluorescent protein and derivatives (Ala206, Leu221, Phe223), and can be suppressed by the mutation A206K. The hydrophobic patch could instead be replaced by a Cysteine-Alanine combination, in which case the self-assembly in nuclei was triggered by addition of a thiol oxidant to the cell culture medium (Bellapadrona and Elbaum, 2016). Such oxidation-induced assemblies were filled rather than hollow and could be seen to sinter, suggesting a liquid-like state.

A connection between protein self-assembly and condensation was drawn by creating a light-induced linker between the ferritin core and an IDP from the fused in sarcoma (FUS) protein in what was named the “Corelet” system (Bracha et al., 2018). Ferritin was fused to an iLID domain via an FP in one color, while a condensing IDR domain was fused via an FP in a second color to SspB. iLID and SspB dimerize under blue light illumination, leading to self-assembly of a network of ferritin proteins linked by interactions between the concentrated IDR. This model system elegantly recapitulates many of the salient features of biomolecular phase separation. Using FPs of distinct colors and careful fluorescence intensity calibrations, the phase space for condensation could be mapped semi-quantitatively.

In this work, we explore the use of optical refractive index (RI) as a diagnostic tool for condensation. Quantitative phase imaging has been used for measurements in vitro without the concerns and constraints associated with fluorescent labels (McCall et al., 2023). The power of RI mapping of cells has also been demonstrated (Kim et al., 2016, 2023; Nygate et al., 2020; Park et al., 2018; Schürmann et al., 2016) by a number of phase-sensitive and holographic microscopy techniques, summarized in a recent review (Kim et al., 2024). Here, we employed the commercial CellExplorer system (NanoLive SA, Switzerland). All measurements were made on live cells, as we reasoned that chemical fixation, whose nature is amino acid crosslinking, risks altering the delicate balance of entropy and interaction that leads to phase separation in the first place. As expected, the condensed phases were more refractive than the dispersed. Less expected were observations that fluorescence intensity could remain high in defined intranuclear spaces even when the RI indicated loss of condensation. Also unexpected were observations of persistent condensation after sufficiently long blue light exposure, even after the blue light was extinguished, and of a further compartmentalization within the dense phase of the Corelets under ferritin-rich conditions. These results suggest that the condensation behavior is richer than the simple biphasic coexistence that is often presumed as a starting point for analysis and also point to a difference between the phase separation reported by fluorescence intensity and the dielectric response of a condensed phase as detected by refractive index.

## Materials and Methods

### PBD IDs used in Fig. 1

**GFP**: 1GFL: https://doi.org/10.2210/pdb1GFL/pdb; **24-mer human ferritin**: 5N27: https://doi.org/10.2210/pdb5N27/pdb; **iLID domain**: 4WF0: https://doi.org/10.2210/pdb4WF0/pdb; **mCHerry**: 6YLM: https://doi.org/10.2210/pdb6YLM/pdb; **SspB**: 1OX9: https://doi.org/10.2210/pdb1OX9/pdb.

**Figure 1.**
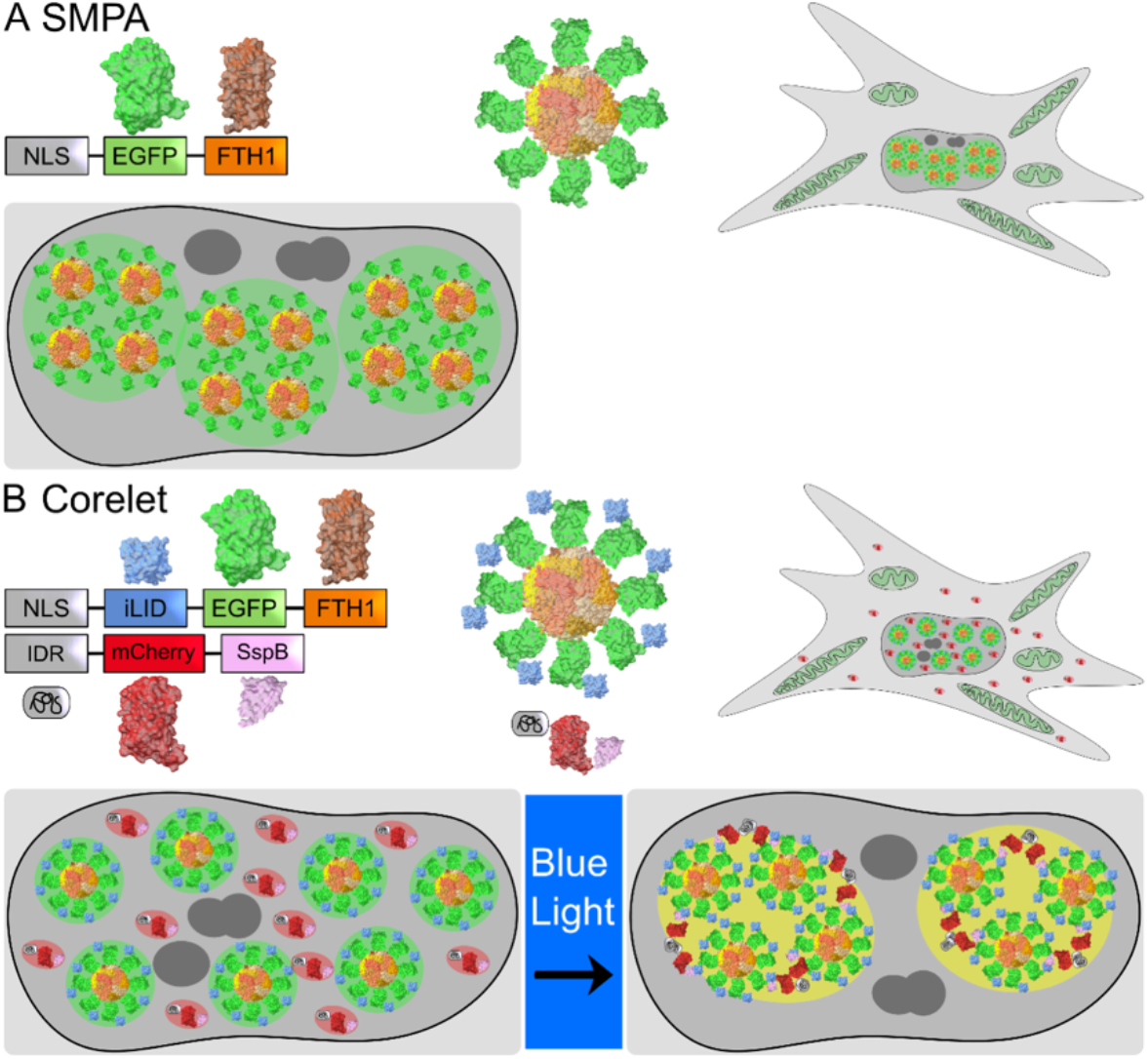
Overview of the Supramolecular protein assemblies (SMPA) and the Corelet system. A) Schematic representation of the SMPA. All structures are taken from the PDB (GFP: 1GFL, Ferritin: 5N27). The assembly of the SMPA is approximated. B) Schematic overview of the Corelet system. PBD structures include GFP (1GFL), Ferritin (5N27), iLID (4WF0), mCherry (6YLM) and SspB (1OX9).

### Plasmid preparation

The one-component SMPA plasmid for NLS-Citrine-ferritin in the pcDNA3 vector was described in a previous publication (Bellapadrona and Elbaum, 2014) and used as is after amplification. The two-component Corelet plasmids for NLS-iLID::EGFRP::FTH1 and FUS_N_::mCherry::SspB (Bracha et al., 2018) were recloned into pcDNA3 from lentiviral vectors received from the Brangwynne lab. Sequence maps are provided in the Supplementary Material. Plasmids were amplified in DH5α cells and purified using the Promega® PureYield™ kit. Plasmid maps are provided in the Supplementary Material and are available upon request.

### Cell culture and transfection

U2-OS and HFF-1 cells were obtained from ATCC. Cell cultures were maintained under 5% CO2 at 37 °C in DMEM medium (Dulbecco®) with 5% fetal calf serum (Biological Industries, Israel).

DNA transfection was performed on cultures at approximately 75-90% confluence using JetOptimus (Polyplus), following the manufacturer’s protocol. SMPA in Fig 2 were induced by transfection with 1.3 µg DNA. Dual transfections at different ratios were performed to induce Corelet formation. DNA stoichiometry Ft:FUS included a roughly 1:3 (Figs 3-5), a high Ft:FUS plasmid ratio to generate larger condensates (5:1 and 10:1 in Figs 6,7, respectively), and a lower ratio to produce small condensates (1:8, Fig 8).

**Figure 2.**
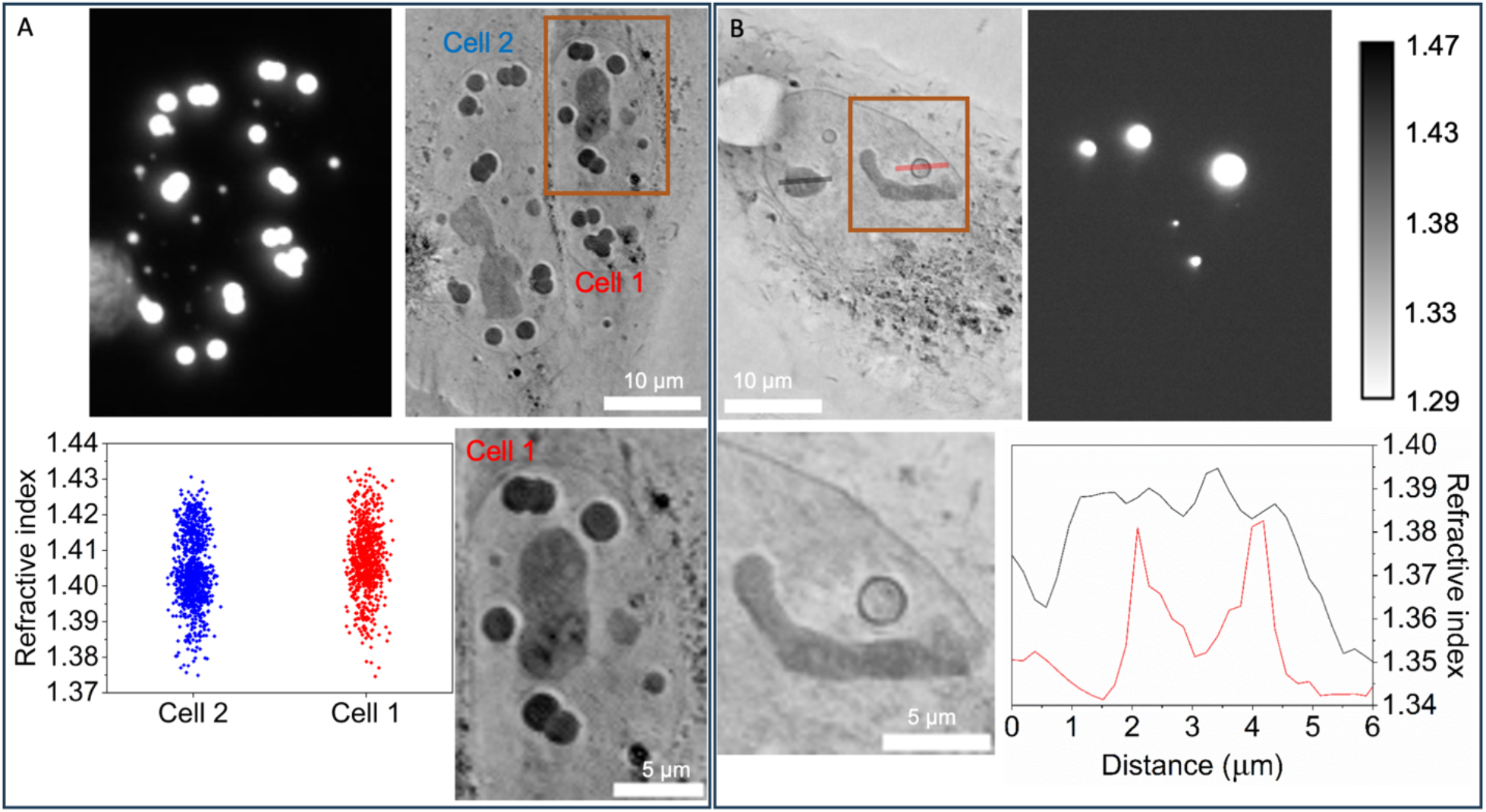
Fluorescence and refractive index measurements of one-component ferritin SMPA. A) Both large and small condensates are seen in the fluorescence, and the corresponding locations show a high RI. Note the clusters of abutting SMPA; they appear to sinter but not fuse, suggesting a solid rather than liquid physical state. Note too, the variety of contrast levels represented, as indicated in the plots, as well as the hints of internal structure seen in the inset. B) SMPA may also take a hollow shell form, seen in the RI even when unresolved in the fluorescence image. Additional examples of SMPA in the cell nucleus of U2OS and HFF cells are seen in Fig S1.

**Figure 3.**
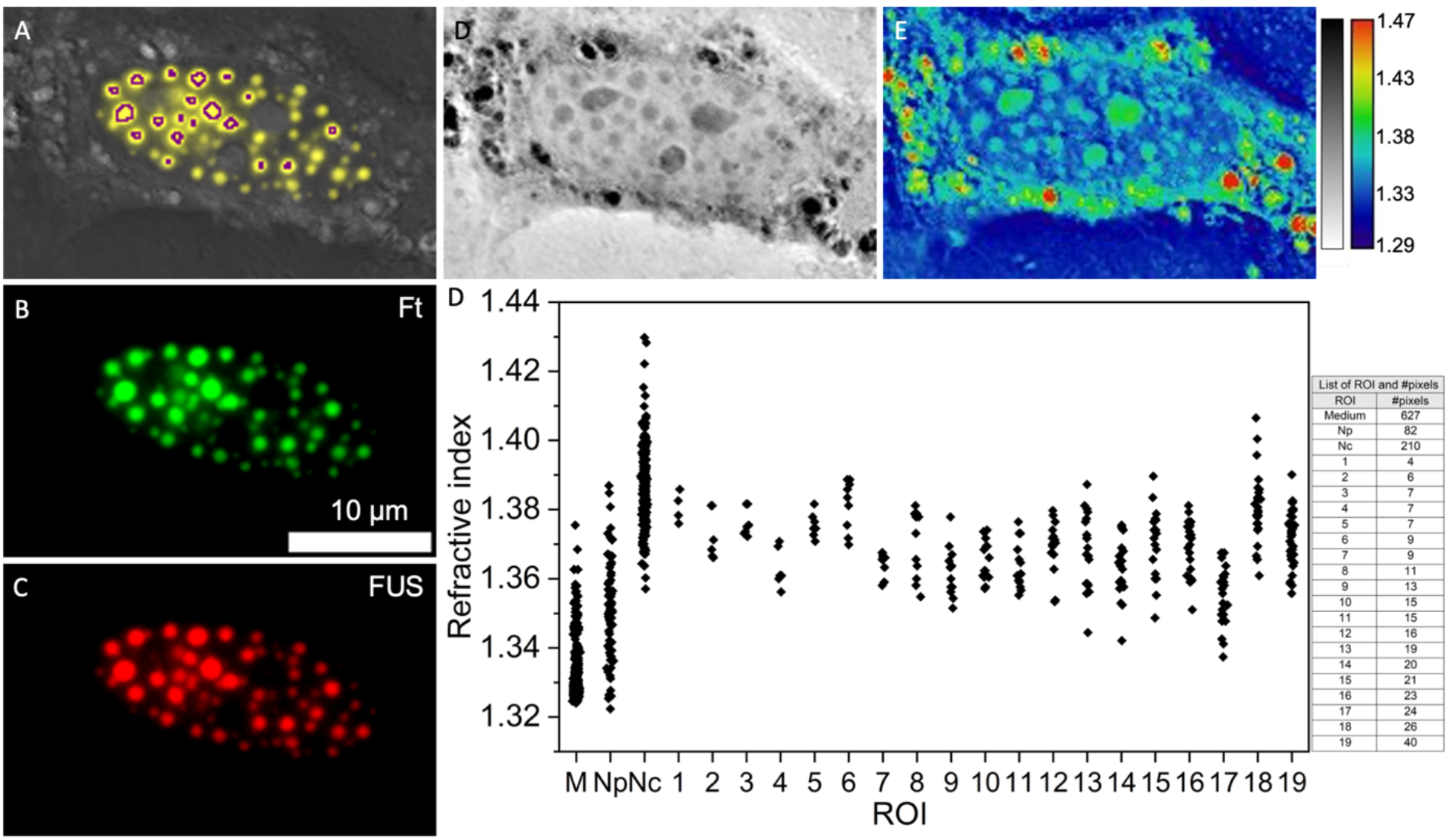
Quantification of RI for two-component Corelet condensates. A) FUS and ferritin (Ft) overlayed on the RI image. The areas of condensation are delineated according to an intensity threshold, eroded morphologically to compensate for fluorescence haze effects, and indicated in purple outline. B) The two fluorescence channels appear in green (Ft) and red (FUS), respectively. C) The RI images are shown in grayscale and false color for enhanced visibility. C) The plot shows a distribution of RI values extracted from the surrounding medium, the nucleoplasm (Np), the nucleoli (Nc), and the 19 regions of interest (ROI) that delineate the Corelet condensates. The associated table shows the number of pixels included in the measurements. Condensates smaller than 4 pixels were removed from the analysis.

**Figure 4.**
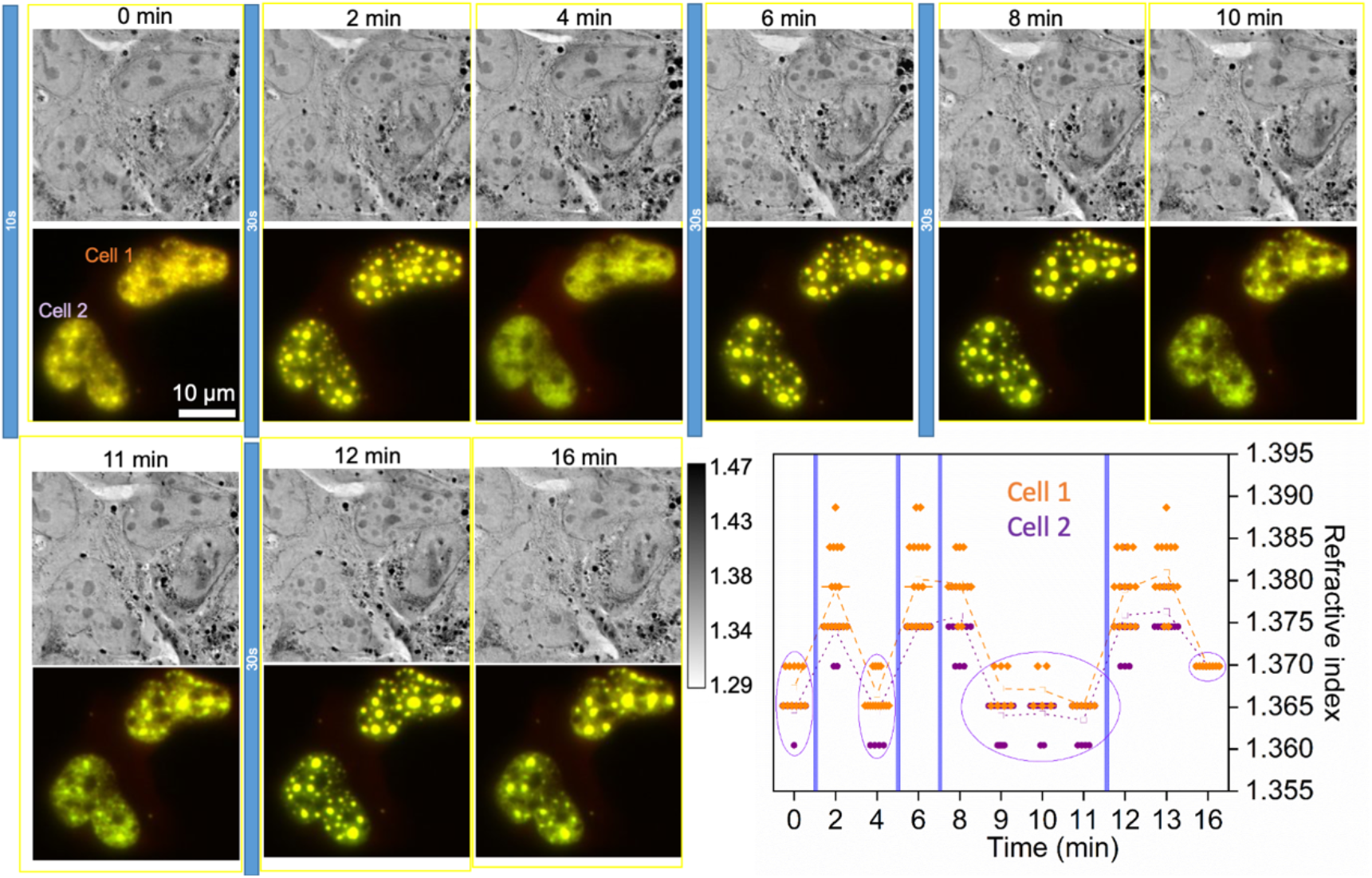
Condensation tracks photoactivation. A) A time series shows that refractive index and fluorescence track the condensation induced by blue light illumination. Illumination periods are indicated by vertical blue bars. With repeated cycles, the fluorescence does not diffuse completely but the RI indicates complete condensation and decondensation. B) A plot of RI measurements according to the method of Fig 3. ROI are defined anew with each condensation and used for the remainder of the cycle to determine RI. RI values have been rounded off at steps of 0.005 for clarity.

**Figure 5.**
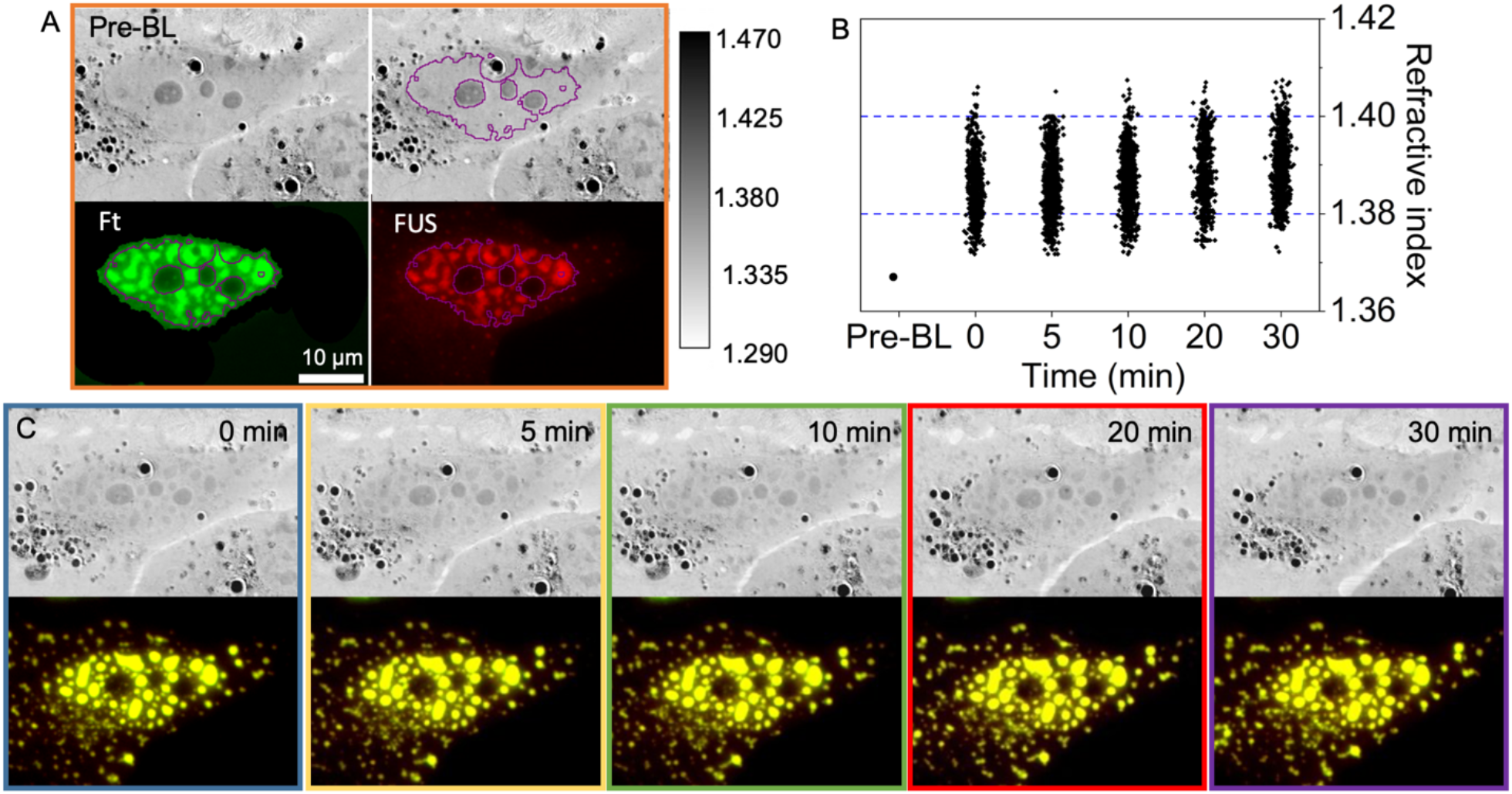
Long activation time induces persistent condensation. A) Prior to blue light exposure (pre-BL) the cell shows no indication of condensation by RI. Fluorescence is distributed unevenly within the nucleus. B) A time trace of images recorded after 20 min continuous activation, showing that the condensates do not disperse. C) Quantification of the RI in condensed regions shows that the RI remains constant for at least 30 min. See Figure S2 for an additional example of long activation resulting in persistent condensation.

**Figure 6.**
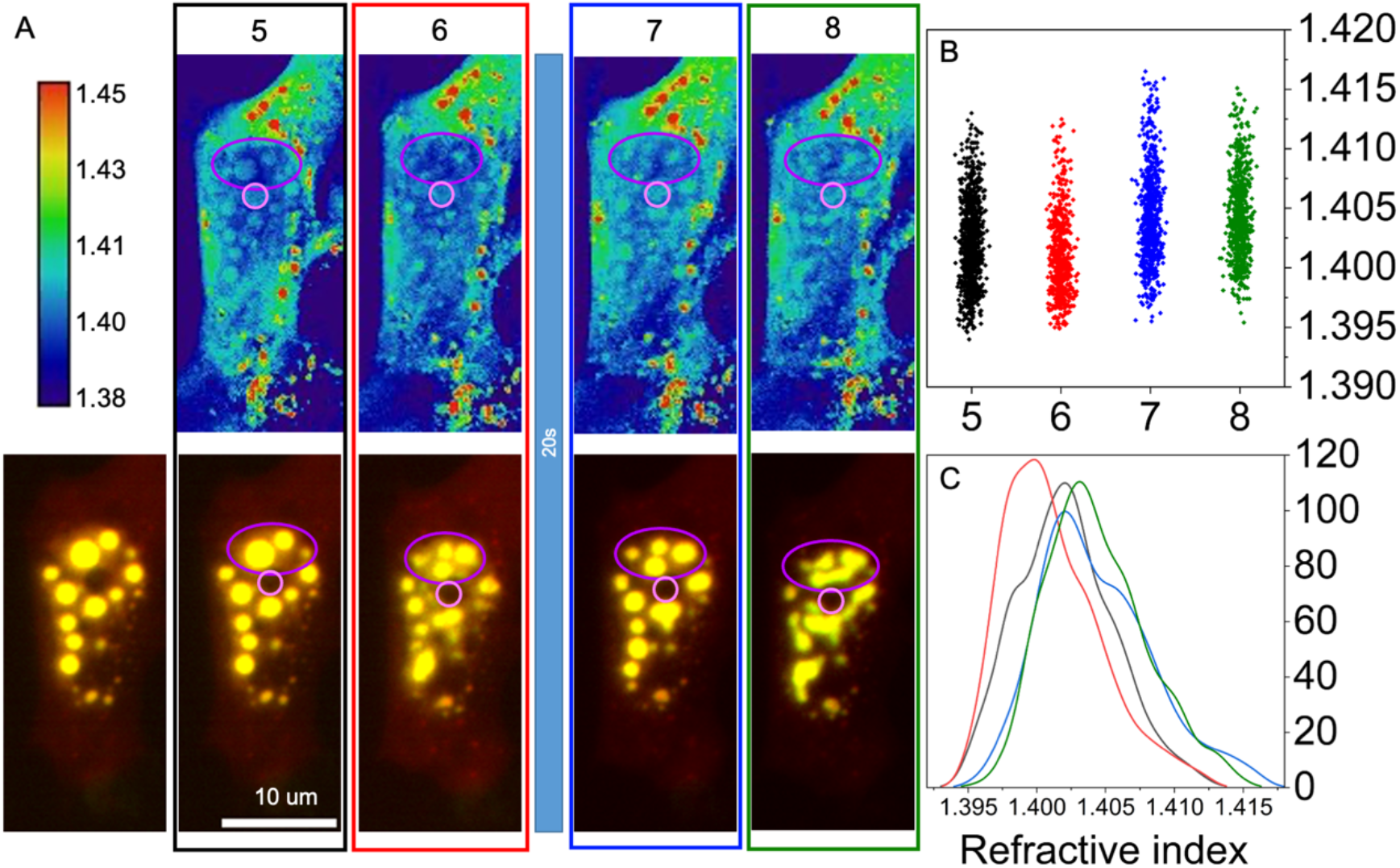
RI indicates marginal condensation persistence. The cells were first illuminated continuously for 10 minutes (not shown) and then kept in darkness (other than rapid illumination for fluorescence imaging with exposure of 1 sec at 30-second intervals) for the number of minutes indicated. A) Decondensation is clear by 6 min in the RI but not in the fluorescence. A subsequent illumination for 20 sec restores the condensation. By 8 min, the condensates are seen to fuse, suggesting the fluid state. Purple and pink circles highlight an example of the decondensation imaged in RI imaging and the nucleoli, respectively. B) Quantification of the RI measurements shows subtle changes during this episode, with a reduction in RI between 5 and 6 min and then restoration at 7 min. This is seen in the points distribution as well as the histogram of RI values, with corresponding colors indicating the time points.

**Figure 7.**
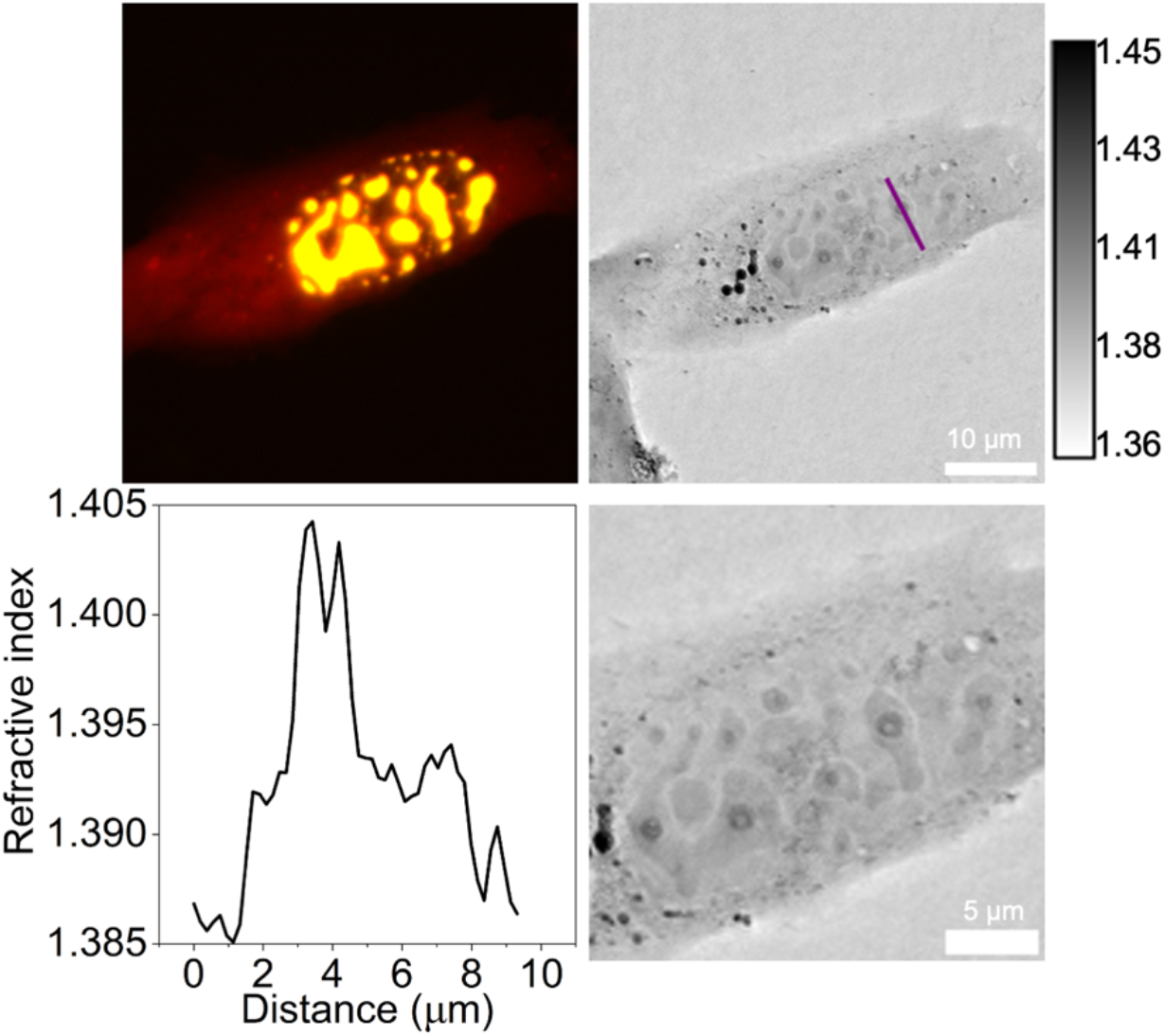
Very large condensates show an internal compartmentalization. A) Fluorescence shows a typical pattern of condensation with high level of protein expression, with non-spherical shapes and apparent fusion. B) The RI images show identical shapes but with much better resolution. A very dense sphere appears within several of the condensates, often with an internal hollow core. The RI of the surrounding condensate is comparable to or slightly less than that of the nucleoli, but the RI of the cores are much higher and recall those of the SMPA seen in Fig 2.

**Figure 8.**
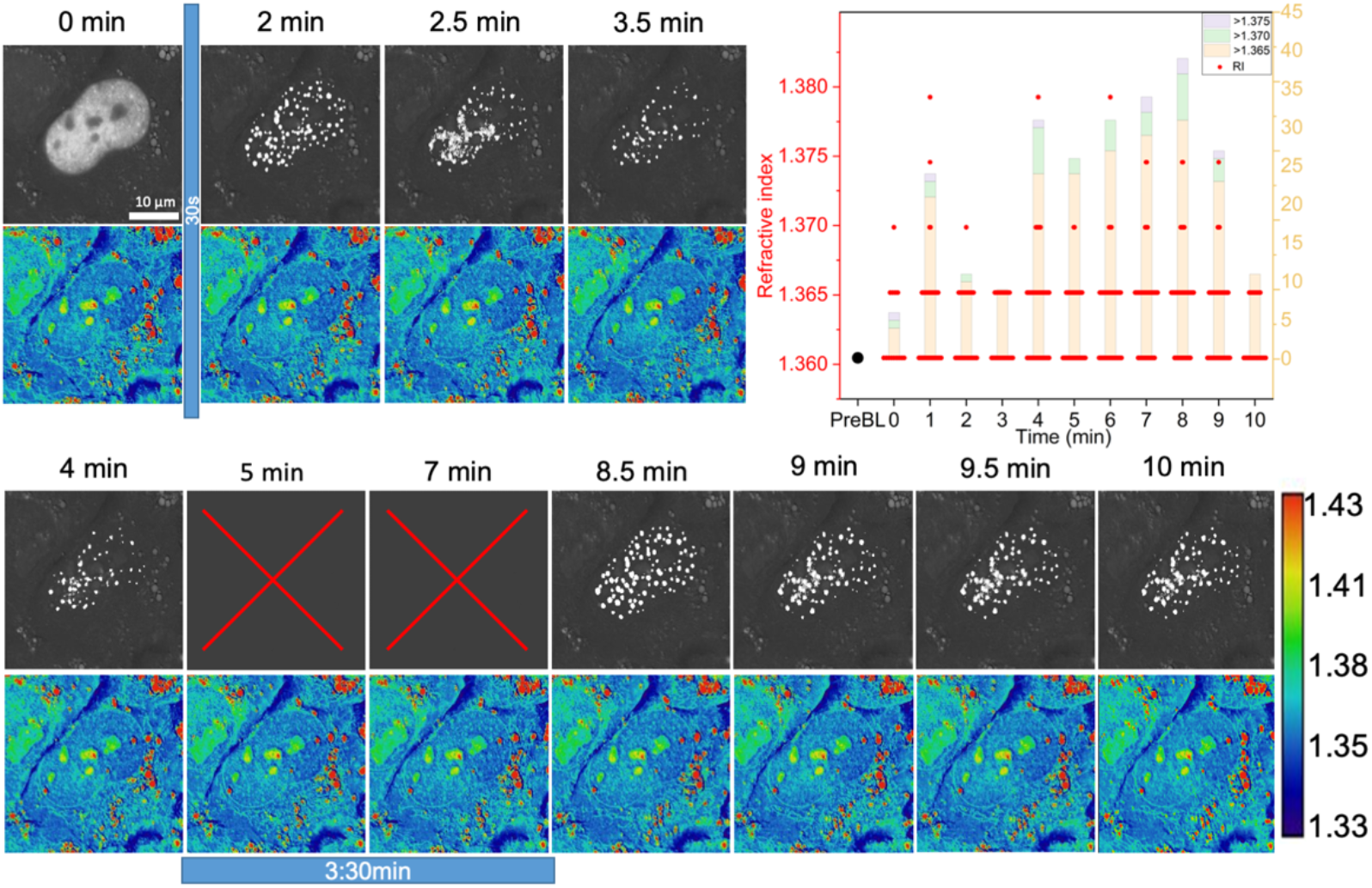
Very small condensates have RI similar to larger ones. A) Condensation induced by 60 sec illumination is seen clearly in the fluorescence. Corresponding points appear in the RI, which peak and then dissipate, but they cover only one or a few pixels. Essentially the same behavior is seen for 30 sec and 210 sec illumination. Supplementary Movie S shows the entire time series at 30 sec intervals; the flashing points in the RI color map are more noticeable there. B) The quantification method of Fig 3 was modified so that all pixels included in condensate ROIs are plotted (with values rounded for clarity), together with bars showing the number of pixels exceeding threshold values. Clearly the RI increase during periods of illumination and decrease following, but this is difficult to follow in the fluorescence alone.

### Refractive Index Measurement

Refractive index mapping was performed in the CellExplorer3D microscope (NanoLive SA, Switzerland). The microscope is equipped with an on-stage incubator to maintain an environment of 5% CO_2_ at 37 °C, as well as fluorescence illumination for green and red emission. The microscope is controlled by the manufacturer’s “Steve” software, which also provides tools for preliminary analysis. The measurement protocol includes a preset self-calibration routine. We found that the reported values were lower than expected and somewhat variable, so we established a second stage of calibration based on RI of polystyrene (Polysciences, 3.0 µm, n=1.600) and silica (Bangs Laboratories, 1.1 µm, n=1.440) nanobeads, and water (n=1.333). This calibration yielded an n=1.45 for the intracellular lipid droplets, which were used as an internal standard.

Photoactivation of the Corelets was performed using a stand-alone LED illuminator with a 22mm aperture, 470 nm wavelength, LED Type A (3W), bare-wire connector (LCS-0470-03-22, Mightex, CA, USA), controlled by an SLA-series two-channel LED Driver (SLA-1000-2, Mightex). The illuminator was mounted diagonally approximately 12 cm from the specimen dish. The LED was focused by a lens to cover an area of 2.6 cm diameter on the dish.

Image processing and downstream ROI analysis were performed using Fiji (Schindelin et al., 2012).

## Results

The protein systems under study are displayed in Fig 1. We will refer to the constitutive single-component system as SMPA (Fig 1A) and the photo-activated, two-component system as Corelets (Fig 1B). Both form in the cell nucleus. SMPAs begin to appear approximately four hours post-transfection, while Corelet components require a somewhat longer expression in order to form condensates upon blue light illumination. Both were examined typically 12-24 hours after transfection. The primary tool in the study is a holo-tomographic microscope (3D Cell Explorer, NanoLive SA, Switzerland), which creates a volume map of the RI using highly tilted illumination recorded in multiple projections around an axis. We found that the RI measurements show high precision and remarkably low noise. Quantification required several calibration steps, as described in the Materials and Methods. The microscope is also equipped for wide-field fluorescence illumination and with an environmentally controlled chamber for extended live-cell observation.

An RI map of the SMPA appears in Fig 2, with two adjacent cells in panel A. The nuclear envelopes are visible in the images. SMPA are identified unambiguously by fluorescence, and contrast in the RI image is very strong. Several clusters of adjacent spheres are seen. Since their growth occurs over many hours, at the rate of protein expression, they appear to sinter as particles but not to fuse as a viscous fluid. The inset also hints to some internal structure that is poorly resolved. This is consistent with the alveolar structure reported in the original work (Bellapadrona and Elbaum, 2014). Nucleoli are also seen in the RI but not in the fluorescence. Notably, the RI of the nucleoli is lower than that of the SMPA. The distribution of RI values could be quantified by using the fluorescence image as a mask overlaid on the RI image. An extreme example of internal structure appears in Fig 2B, where the SMPA appear in the RI image as thin spherical shells. The fluorescence signal was easily saturated, on the other hand, due to limited dynamic range of the 8-bit recording, and the central hole does not appear. Saturated fluorescence is a common problem in the field, as a large dynamic range is required in order to display both the bright condensate and the dim dilute phase. Moreover, out-of-focus fluorescence will make the condensates appear larger than they truly are. A line profile across the hollow sphere shows the high RI concentrated at the SMPA rim. Other examples appear in Supplementary Fig S1. Given the dimensions of a single voxel, it is likely that the RI measurement is somewhat underestimated due to cross-coverage with the surrounding nucleoplasm.

Preliminary observations of Corelets showed a similar correlation of fluorescence with RI, but analysis was more challenging due to the lower RI contrast and smaller size of the condensates. Fig 3 establishes the visualization and analysis strategies. It displays the fluorescence in the two individual colors, green and red, for ferritin (Ft) and FUS, respectively, as well as a composite image in yellow. A threshold imposed on the fluorescence intensity delineated the regions of high protein concentration as regions of interest (ROI) in ImageJ, shown outlined in purple. The ROI boundaries were eroded morphologically in order to compensate for the spread in the fluorescence signal that originates from planes out of focus. (Erosion by 0, 1, or 2 pixels was done on a case-by-case basis.) These ROI were then used as selections for the corresponding image locations in the RI map. Note that the RI map is volumetric and resolved in depth. A single 2D slice is used for display with an inverted lookup table (higher index dark), as well as a blue-red (“physics”) color scale. For quantification, several slices are chosen so as to limit to regions within the nucleus, and a maximum intensity projection is used in order to match the volume from which the fluorescence emerges. (The method is prone to error when more refractive elements, such as nucleoli, encroach into the contour-defined area in planes above or below the condensed body of interest. In such cases, the measurements must be discarded.) Corelet condensates are always more refractive than the nucleoplasm and chromatin background, but unlike the SMPA, they are typically less refractive than the nucleoli.

With the quantification protocol in hand, we first recapitulate the dynamic behavior of the Corelet system. Fig 4 shows a cycle of condensation, dispersal, and re-condensation upon transient blue light illumination. A rapid illumination (10 sec) was used first to find the transfected cells in which condensation occurs. Subsequent cycles with 30 sec illumination showed a rapid response of the RI signal, whereas the fluorescence showed a hysteresis with the signal intensity, remaining elevated nearby the former condensate locations. The condensation likely created 3D voids in the chromatin, away from which diffusion would be slow. After repeated cycles, the persistent fluorescence became more clearly defined. (Compare 4 and 16 min time points.) Fluorescence intensity reveals the local protein concentration. Refractive index, on the other hand, relates to dielectric polarizability. It reflects the bulk material continuum rather than the concentration of isolated molecules. Thus, the continuum, condensed state is rapidly lost even though the local protein concentration remains high.

Unexpectedly, after an extended illumination of 20 min with blue light, the condensates did not disperse over a period of more than 30 min (Fig 5). Fluorescence remained punctate and intense, and the RI remained uniformly high at the same locations. This indicates an annealing process and suggests that the internal structure of the rapidly responding condensates differs from that of the annealed ones. A possibility to consider is the transition to a gel state. However, the FUS moiety is interspersed with the ferritin. Given the two-component assembly mechanism, one may expect that the internal FUS concentration may reconfigure in a manner that imparts stability to the composite assembly. In order to test the stability, we illuminated for a shorter extended period of 10 min (Fig 6). After 6 min in darkness, the Corelet condensates began to disperse according to the RI contrast. Quantification showed a skew of the RI distribution to lower values, as seen in the histogram. The fluorescence still indicated a condensed state, albeit with a subtly different distribution. A 20-second re-illumination restored the condensed state at the 7-minute time point, as seen both in the image and in the histogram of RI values. At 8 min, the image reveals the sintering of nearby spheres into elongated shapes; the histogram shows a rearrangement of values at the peak but a remarkable overlap at the upper tail. The flow observed suggests that the dark-persistent condensates do remain in the state of a viscous liquid.

A striking feature is observed in Fig 7: within the very large condensates, a smaller and denser sphere may appear. The RI of the dense sphere is very similar to that of the one-component SMPA, suggesting an interpretation as a ferritin-dense core and ferritin-poor shell surrounding it. At the opposite extreme, an example of very small condensates appears in Fig 8. This presented a challenge for the analytical workflow. Due to the optical effects (diffraction limit and defocus), as well as camera saturation, the size represented in the fluorescence image is much larger than the true size in the sample. Therefore, the mask defined by the fluorescence over-estimates the relevant area severely. In order to detect such condensation, we present all the RI voxel values within the fluorescence-defined mask and focus attention only to those few pixels taking values measurably above the background. These high-valued pixels, indeed, responded to blue light illumination as expected for repeated short exposures. In the RI images, these outliers are difficult to resolve by eye in gray levels; they can be seen more easily on the color scale. Compare, for example, the 8 and 10 min points, where the Corelet condensates appear almost as noise in the earlier measurement but can be identified clearly by correlation with the fluorescence. The peak values fall in the range described in Fig 4, close to 1.380 and so the internal structure of the very small condensates is likely similar to that of the larger ones.

## Discussion

Protein condensation in cells, and particularly the phenomenon of liquid-liquid phase condensation, has attracted enormous attention as a mechanism for biochemical regulation in vivo. The biological examples are broad and varied. Synthetic systems offer a more controlled platform for investigating basic biophysical principles. In this light, the two ferritin-based systems offer a bridge between the concepts of kinetic self-assembly and the thermodynamics of phase condensation. While simple theoretical considerations posit coexistence between individually sparse and dense phases, the observations here indicate an evolution of the dense phase over time. Specifically, illumination for 20 minutes or longer resulted in persistent condensation lasting more than 30 minutes in the dark. Illumination for 10 minutes resulted in condensation lasting approximately 5 minutes, with slow evolution of the shapes that suggests retention of the viscous liquid-like physical state. Illumination for 30-60 seconds confirmed the reversibility reported previously. We may, therefore, ask what internal changes occur in converting the transient to the persistent state.

In the one-component SMPA, each ferritin sub-unit is hybridized to a dimerizing unit in the FP. Thus, each ferritin protein holds 24 potential linkers. The resulting SMPA is stable and solid-phase. The high density is reflected in the high RI values, equal to or typically greater than those of the nucleoli. The Corelet condensates show a lower RI than the SMPA, typically lower even than that of the nucleoli. RI values also vary more significantly from one condensate to another, even within the same cell. This may reflect a heterogeneity in the local ferritin:FUS stoichiometry. This will depend on transfection DNA concentration, transfection efficiency, and levels of protein translation. Only the first is under direct control, and the results shown here appear to explore the limits of the phase coexistence boundaries (Bracha et al., 2018).

The dark-persistent condensation upon long activation was unexpected. The RI of long-illuminated, persistent Corelet condensates was not significantly higher than that of the transient ones. Persistent condensation was described almost anecdotally in the original Corelet reference (Bracha et al., 2018), where it was noted that high blue laser intensity in the context of photobleaching might damage the iLID domain of the photoactivatable linker. In the present observations, the persistence resulted from a long activation duration, but the same intensity was not sufficient to cause noticeable photobleaching in any of the experiments. Another suggestion might be to invoke the tendency of the FUS IDR domain to transform to a gel state upon aging (Patel et al., 2015). Irreversible aggregation was indeed observed for the same FUS protein domain in a one-component photoactivatable predecessor to the Corelet system (Shin et al., 2017); these aggregates retained their irregular shapes upon sintering, counter to expectations for a liquid phase. In the present case, the FUS moieties would be interspersed between ferritin cores. A more stable, gel-like interaction between the neighboring FUS might nonetheless form during the long photoactivation, leading to a solid cast of FUS surrounding the ferritin. In the subsequent period of darkness, the equilibrium affinity of the iLID-SspB interaction should drop drastically, but the contact may not actually rupture if the two components are held together externally. For the 20 min photoactivation, the condensates were stable for at least 30 minutes, whereas for the 10 min photoactivation, a partial decondensation was detected after 7 min. Recondensation after a short blue-light exposure shows that the dark-persistence is not due to inactivation of the iLID. It is also unlikely that the FUS domain reached an irreversible prion state on this time scale, especially given the maintenance of a spherical shape and the absence of any detectable change in RI.

We consider next the use of RI as a diagnostic for protein condensation. The measurement is compatible with live imaging and provides very useful depth resolution without the complications of fluorescence or confocal imaging, as well as lower risk of phototoxicity. Also, while fluorescence may be required for molecular identification, there is always a risk of interaction induced by the FPs themselves. We see that the RI responds specifically to the condensed state, whereas the fluorescence measures, in practice imperfectly, only the local protein concentration. As is apparent from Fig 4, repeated cycles of Corelet condensation and decondensation in the nucleus create a space from which diffusion is slow. Presumably, this is a void that forms within the chromatin; this would be consistent with previous reports on protein condensate interaction with chromatin (Shin et al., 2018). The boundaries of fluorescent puncta cycle between sharp and blurred, yet the high local concentration remains. Compare, for example, time points 12 and 16 min, or 8 and 10 min. Nonetheless, the RI responds immediately to the transient illumination and darkness. Upon de-condensation, no visible trace is left in the RI image. We cannot rule out the possibility that some oligomeric form of the Corelets remains intact, perhaps slowing the diffusion process due to larger hydrodynamic diameter. Upon longer activation, as seen for example in Fig 5 and Supplementary Fig S2, the condensates are stable and both RI and fluorescence signals appear constant. With an activation of intermediate duration, seen in Fig 6, the RI proves more responsive and reliable as a diagnostic of Corelet condensation. The RI can also detect structure within the condensate, for example the dense cores in Fig 7 or the hollow centers for the SMPA in Fig 2, where there is also a hint of inhomogeneity within the densely filled bodies. For very small condensates, the RI puncta covering only a single or very few pixels are very difficult to identify from noise without the fluorescence signal, and yet the fluorescence alone is not a reliable indicator of their state of condensation.

The relation of protein concentration in solution and refractive index has been analyzed in the context of a mixing model based on independent contributions of the constituent amino acids (Kassimi and Thakkar, 2009; Zhao et al., 2011). This relation is described as a refractive index increment, or derivative dn/dc, and the mass density or concentration in solution is extracted from the linear proportionality. The model has been extended to consider hydration shells and protein structure (Khago et al., 2018), as well as more complex molecular composition in solution (Beck et al., 2024; Möckel et al., 2024). Here we observe a dramatic effect on RI of the state of condensation, i.e., the transformation of isolated macromolecules into a material continuum, without a very dramatic change in concentration of the condensing proteins. The refractive index is a measure of dielectric polarizability at the optical frequency. Modeling of such polarization as an additive contribution of isolated dipoles is akin to a noble gas approximation, which ignores the possibility of extended electronic excitation and correlation at longer length scales. Thus, the molecular connectivity and linkage in self-assembly is likely to play a crucial role, and the extrapolation of mass density from RI measurements of condensates, rather than solution, should be approached with some caution.

Another recent work reported on RI measurements of phase separation of nucleoli, as well as heterochromatin, nuclear speckles, and cytoplasmic stress granules (Kim et al., 2023). Our observations of nucleoli as dense condensates are very similar, and we too saw no hint of nuclear speckles or other nuclear protein condensates. (We did not induce or examine for cytoplasmic stress granules.) Quantitatively, our numerical measurements for the nucleoli are somewhat higher. This may reflect a difference in calibration protocol or the measurement technology, as our numbers for lipid droplets are also higher than a previous report (Kim et al., 2016). We consider this a minor discrepancy, however, since we do not aim to quantify mass density, and we use the nucleoli and lipid droplets as an internal standard against which to compare the synthetic protein condensates. Notably, the one-component ferritin SMPA, in which the hybridized linkages may saturate in close packing, showed a much higher RI than the Corelet condensates, where the linkers are more sparse. The RI of SMPA were equal or significantly higher than RI of the nucleoli in the same cells. A comparison with the low-density condensates, *viz*., speckles and stress granules (Kim et al., 2023), is perhaps even more interesting. These have been defined and studied exhaustively by means of fluorescence imaging but do not present any measurable signal in the RI. At the same time, it was shown that they remain permeable to diffusing fluorescence protein probes, indicating an open, porous structure. This is consistent, then, with the notion of a minimal protein-protein connectivity required for the dielectric polarizability to reflect a material continuum. Thus, the condensed state reported by RI is subtly different from the notion of thermodynamic phase separation per se. As had been pointed out (Kim et al., 2023), phase separation may result, for example, from depletion interaction or polymer segregation rather than associative interaction, or from sparse and possibly labile cross-links between long polymers such as RNA. Such low-density condensates may appear in fluorescence without forming a polarizable dielectric continuum distinct from the solvent.

The simple two-phase coexistence model for phase separation posits a uniform concentration within each of the high and low density regions. Clearly, in the example of Fig 7, the condensed phase is not uniform, and a certain phase separation occurs within. This complicates the simple picture of two-phase equilibrium, but may offer an experimental paradigm for layering or compartmentalization of more complex mixtures. Such multi-component systems have been addressed theoretically (Jacobs and Frenkel, 2017), and recall the core-shell structure reported for nucleoli on the basis of fluorescence observations (Brangwynne et al., 2011). Although it was not the aim of the present study, we may also point out that the RI contrast of nucleoli suggests some internal compartmentalization.

## Conclusion

In summary, we show that refractive index mapping is a useful addition to the toolchest for study of protein condensation and liquid-liquid phase separation. It is especially useful for study of live cells where photo-damage is a major concern. In comparison with fluorescence imaging, it is quantifiable and much less subject to artifacts such as saturation (high or low), or effects of out-of-focus sources. Combination of 3D RI mapping with high dynamic-range confocal imaging would offer a further step forward for analysis. Using synthetic ferritin-based condensates expressed in living cells, we have shown that the RI reveals a transformation from an elevated local concentration to a truly condensed material phase with an elevated optical polarizability. This should provide further clarification and classification of biomolecular condensates and their condensation dynamics.

## Acknowledgments

The authors acknowledge the lab of Cliff Brangwynne for provision of Corelet plasmid sources, with special thanks to Dan Bracha for discussions and comments. In addition, the authors acknowledge the help of Joseph Addadi, of the Life Science Core Facilities, for the help with the microscopic acquisition. We acknowledge the de Picciotto Cancer Cell Observatory, In Memory of Wolfgang and Ruth Lesser, in which the CellExplorer3D microscope was operated. This work was funded in part by the US-Israel Binational Science Foundation and by the European Union (ERC AdG, CryoSTEM, 101055413. Views and opinions expressed are however those of the authors only and do not necessarily reflect those of the European Union or the European Research Council. Neither the European Union nor the granting authority can be held responsible for them.) ME is incumbent of the Sam and Ayala Zacks Professorial Chair in Chemistry. The Elbaum lab has benefited from the historical generosity of the Harold Perlman family.

## Notes

### Competing Interest Statement

The authors have declared no competing interest.

