## Supplemental Figures for "Refractive index as an indicator for dynamic protein condensation in cell nuclei"

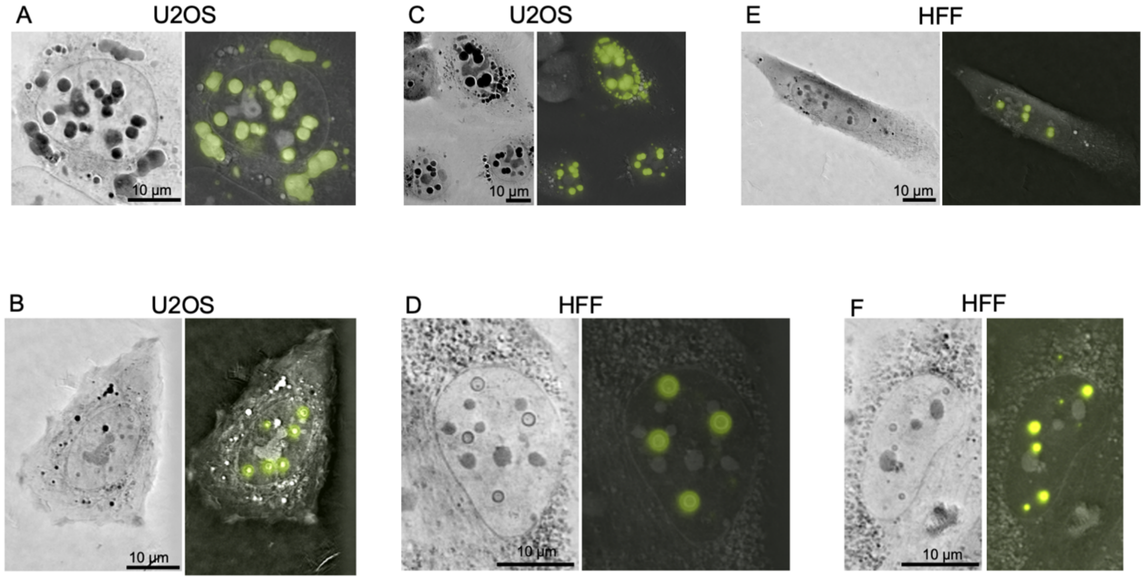


**Figure S1. Additional Examples of SMPA in U2OS and HFF cells.** SMPA can be reproduced in U2OS (A,B,C) and HFF cells (D,E,F). The thin spherical shells are seen in both cell types (B, D, F).

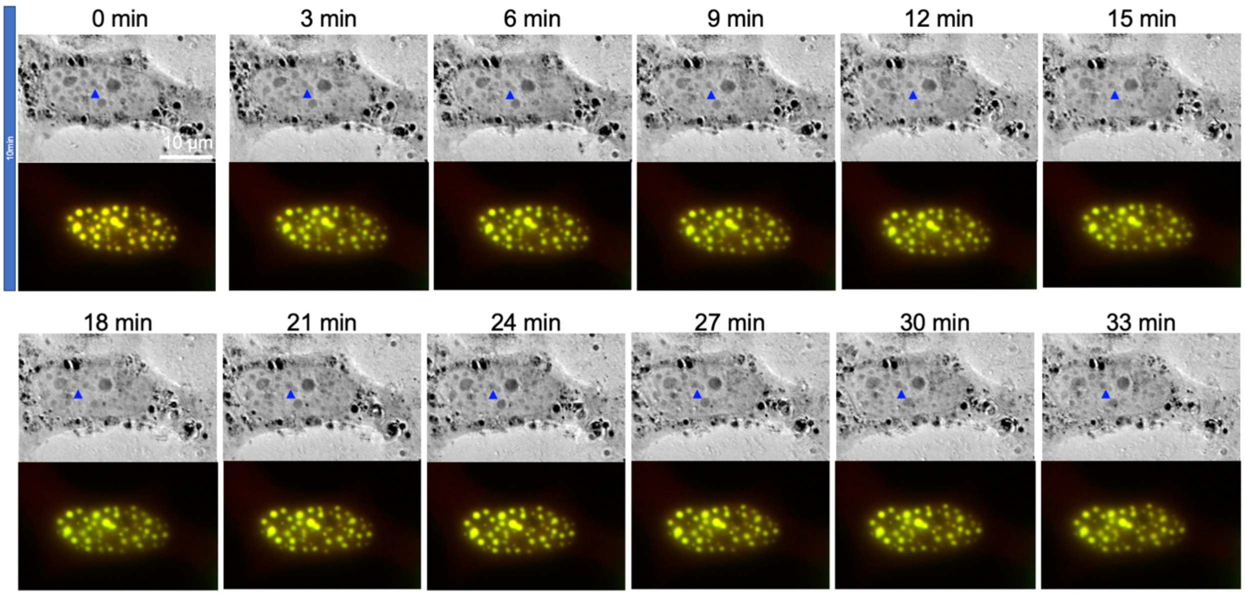


**Figure S2: Long observation of Corelets.** After 10 min blue light exposure, the same cell was imaged for 33 min (1x per minute); every 3 min is depicted. The blue arrowhead points to an example of a persistent Corelet.

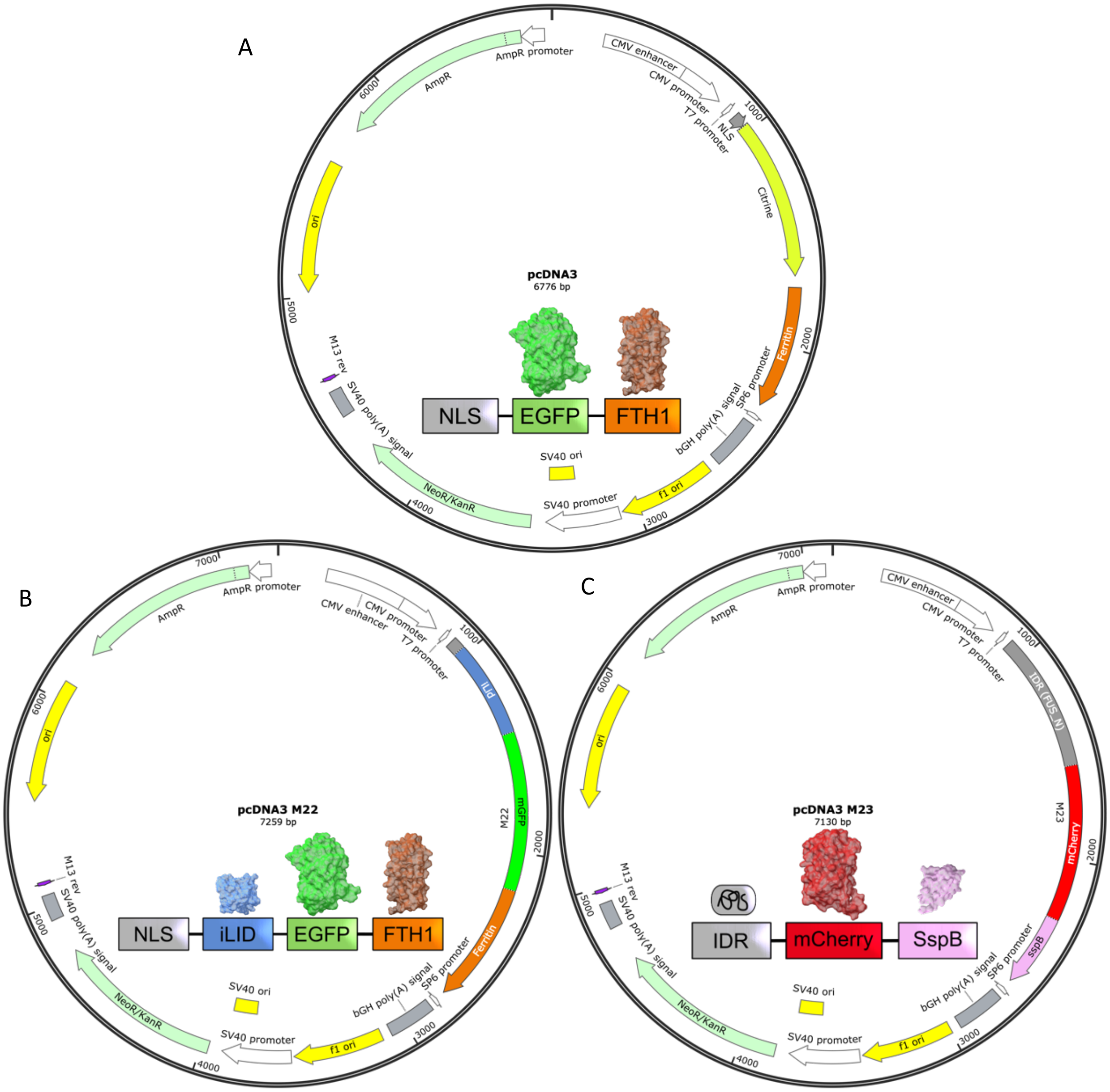


**Figure S3: Plasmid maps of (A) SMLP, (B) the first and (C) second part of the Correlet system in the pcDNA3 vector:** (A) NLS-Citrine-ferritin, (B) NLS-iLid-mGFP-Ferritin, and (C) IDR region of FUS_N-mCherry-SspB.
